# *In vivo Firre* and *Dxz4* deletion elucidates roles for autosomal gene regulation

**DOI:** 10.1101/612440

**Authors:** Daniel Andergassen, Zachary D. Smith, Jordan P. Lewandowski, Chiara Gerhardinger, Alexander Meissner, John L. Rinn

**Affiliations:** Harvard University, Department of Stem Cell and Regenerative Biology, United States; Max Planck Institute for Molecular Genetics, Department of Genome Regulation, Germany; University of Colorado at Boulder, Department of Biochemistry, United States

## Abstract

Recent evidence has determined that the conserved X chromosome “mega-structures” controlled by the *Firre* and *Dxz4* alleles are not required for X chromosome inactivation (XCI) in cell lines. Here we determined the *in vivo* contribution of these alleles by generating mice carrying a single or double deletion of *Firre* and *Dxz4*. We found that these mutants are viable, fertile and show no defect in random or imprinted XCI. However, the lack of these elements results in many dysregulated genes on autosomes in an organ-specific manner. By comparing the dysregulated genes between the single and double deletion, we identified superloop, megadomain, and *Firre* locus dependent gene sets. The largest transcriptional effect was observed in all strains lacking the *Firre* locus, indicating that this locus is the main driver for these autosomal expression signatures. Collectively, these findings suggest that these X-linked loci are involved in autosomal gene regulation rather than XCI biology.

## Introduction

In female mammals, one of the two X chromosomes is inactivated to compensate for gene dosage between males and females ^1^, a process termed X-chromosome inactivation (XCI). Imprinted XCI starts early in development by specifically inactivating the paternal X chromosome (Xp) ^2,3^. While the Xp remains silenced in extra-embryonic linages ^4^, reactivation occurs in the embryo during implantation, followed by random XCI ^5^. After this decision has been made, the inactive X (Xi) chromosome is epigenetically maintained silenced throughout cell division as a compact chromatin structure known as the “Barr body” ^6^.

Recent studies using chromosome conformation capture (3C) based methods have identified that the Xi folds into two “megadomains” and forms a network of long-range interactions termed “superloops” that are directed by the non-coding loci *Firre* and *Dxz4=* ^7,8^. Deletion of *Dxz4* in cell lines leads to the loss of both megadomain and superloop ^9–12^, while deletion of *Firre* alone disrupts the superloop but has no impact on megadomain formation ^11,13^. Moreover, the *Firre* locus is transcribed into the long non-coding RNA (lncRNA) *Firre* that escapes XCI and plays a role in nuclear organization ^14–16^. Recent studies in human and mouse deleted these elements in cell line models of random XCI and found minimal impact on X chromosome biology beyond the loss of these structures ^9–13^.

To date, the phenotypic consequences of deleting these elements for mammalian development has not been addressed. Here, we answer this question by generating mice lacking *Firre* and *Dxz4* and performing an extensive transcriptomic analysis in embryonic, extraembryonic and adult organs. We determined that these elements are dispensable for mouse development and XCI biology. However, the absence of these loci results in organ-specific expression changes on autosomes, suggesting that these regions are involved in autosomal gene regulation rather than XCI biology.

## Results

In order to test the *in vivo* role of *Firre* and *Dxz4* loci both individually and in combination, we generated three knockout (KO) mouse strains: two carrying a single locus deletion (SKO) of either *Firre* ^17^ or *Dxz4* and one carrying a double locus deletion (DKO) of *Firre* in conjunction with *Dxz4* (Fig. 1a, Supplementary Fig. 1a, Material and methods). Notably, we targeted the exact same regions that have been previously reported to disrupt the superloop ^13^ and megadomain structures ^10^. All founder mice were screened by PCR using primers that span the deleted region and identified mutants were confirmed by Sanger sequencing (Supplementary Fig. 1b-c, Material and methods).

**Figure 1.**
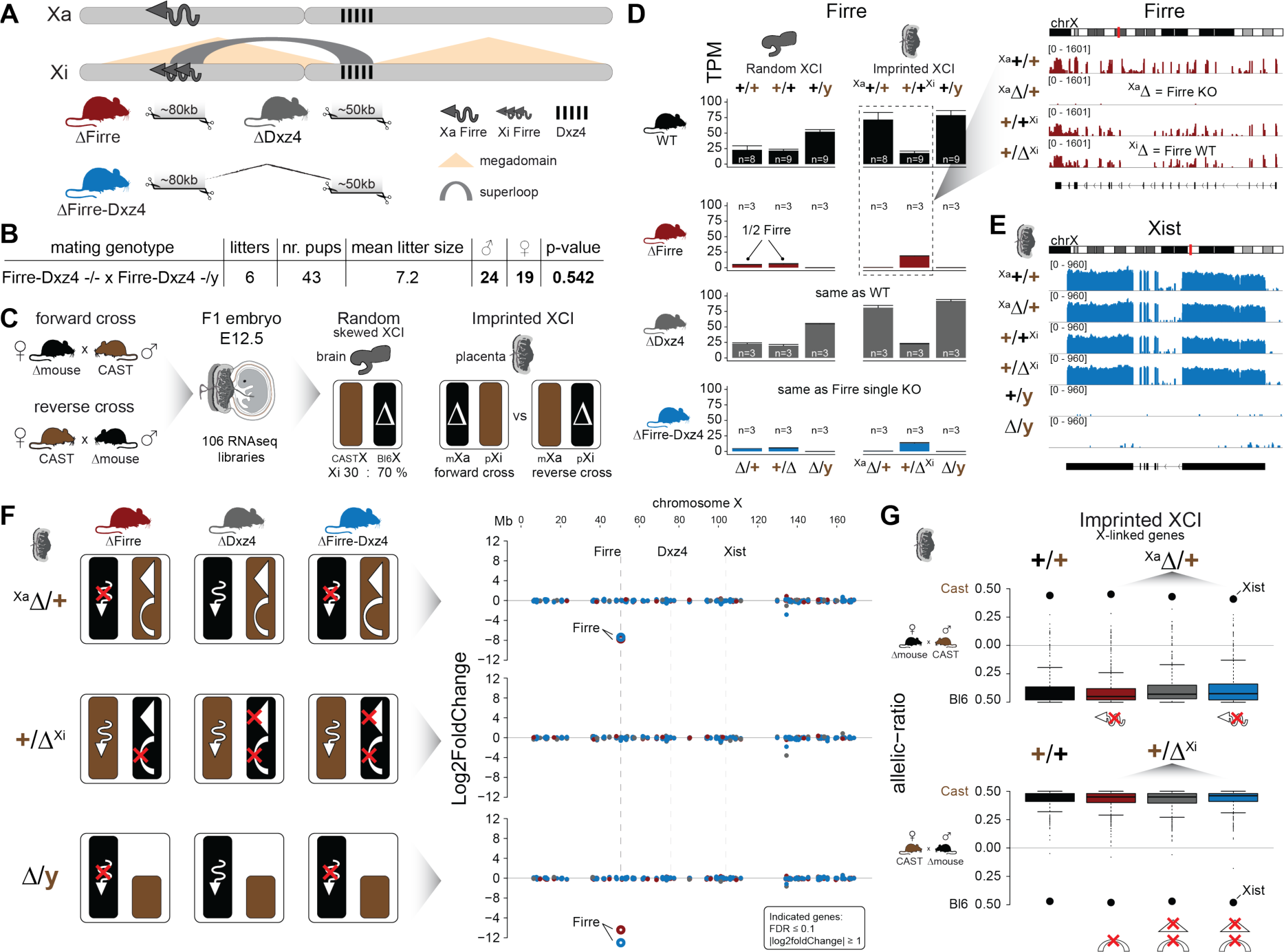
Mice carrying a single or double deletion of *Firre* and *Dxz4* are viable and undergo normal random or imprinted X chromosome inactivation (XCI). (a) Schematic representation of the active (Xa) and inactive X (Xi) chromosomes. The deleted loci of the SKO and DKO mouse strains are indicated (*Firre* (red), *Dxz4* (grey) and *Firre*-*Dxz4* (blue)). The *Firre* locus escapes random XCI and resulting in a full length transcript from Xa chromosome and multiple short isoforms from Xi. (b) Sex type results from homozygote intercrosses. The *P*-value was calculated based on binomial distribution. (c) Allele-specific RNAseq approach to test the functional impact of the maternal and paternal deletion on random and imprinted XCI using the Allelome.PRO pipeline 18. While the brain undergoes random/skewed XCI (Bl6 X chromosome inactive in 70% of cells), the placenta undergoes imprinted XCI (paternal X chromosome 100% inactive). Allele-specific analysis of the placenta allows to distinguish the effects of maternal inheritance of the deletion (Xa, forward cross) versus paternal inheritance of the deletion (Xi, reverse cross). (d) *Firre* expression abundance (mean and SD) in female and male brains and placentas for WT and KO mouse strain. Notably, *Firre* is approximately 4 times higher expressed from the Bl6 allele compared to the CAST, which was observed by comparing the expression levels between the forward cross (Xa Bl6) and reverse cross (Xa CAST) in the placenta. (e) *Xist* expression abundance in placenta for the *Firre*-*Dxz4* double KO strain. (f) Schematic overview showing the effect in the placenta of the deletion on Xa (top), Xi (middle) or in males (bottom) for every KO strain (left). Log2foldchanges across the X chromosome between wildtype and KO strains (right). *Firre* is the only differentially expressed gene on the X chromosome (DEseq2: FDR ≤ 0.1, |log2foldChange| ≥ 1). (g) Boxplot showing the allelic ratios for X-linked genes in the placenta in WT and in the three KO strains, for the forward cross (deletions on the maternal X = Xa) and reverse cross (deletions on the paternal X = Xi).

We found that homozygous mice of all three strains are viable and fertile, and by crossing males and females carrying a homozygous deletion we observed the expected litter sizes and sex ratios (Fig. 1b). To test whether the absence of the *Firre* and *Dxz4* loci has an impact on random or imprinted (Xp chromosome inactive) XCI *in vivo*, we collected embryonic day 12.5 female F1 brains (random/skewed XCI) and placentas (imprinted XCI) from reciprocal crosses between our KO strains and *Mus musculus castaneus* (CAST), followed by RNA sequencing (RNAseq) and allele-specific analysis using the Allelome.PRO pipeline ^18^(Fig. 1c). We further collected the same organs from males to test the role for these loci on the X chromosome outside of XCI biology. Unsupervised clustering of the sequenced samples confirmed the identity of the tissue and the genotype of *Firre* and *Dxz4* (Fig. 1d, Supplementary Fig. 2a-b).

We then investigated the *Firre* expression pattern between wildtype and *Firre* SKO in female brains, and detected approximately half of the wildtype levels as expected for random XCI (Fig. 1d, Supplementary Fig. 2b). Next, we inspected the *Firre* expression pattern in the placenta, an organ that undergoes imprinted XCI, to distinguish between having the deletions on the Xa (maternal inheritance of the deletion) versus the Xi (paternal inheritance of the deletion) chromosome. In contrast to previous reports that identified *Firre* as a gene that escapes XCI in cell lines that model random XCI ^14–16^, we found that *Firre* is exclusively expressed from the Xa, as we did not detect *Firre* expression if the deletion was inherited maternally. Alternatively, we detected wildtype expression levels in the paternal deletion, which confirms that *Firre* does not escape imprinted XCI (Fig. 1d, Supplementary Fig. 2b). Remarkably, we detected the same *Firre* pattern in the absence of *Dxz4*, suggesting that disruption of the superloop between the two loci or the absence of megadomains has no impact on *Firre* gene expression in either the random (brain) or imprinted (placenta) inactivation regimes (Fig. 1d). We did not observe changes in *Xist* expression levels in the absence of these loci (Fig. 1e).

Since *Firre* is only expressed from the Xa in the placenta (imprinted XCI), we can disentangle the functional role of *Firre* RNA from megadomain and superloop structures that only exist on the Xi. We hypothesized that: (1) females and males carrying a maternal *Firre* single or *Firre*-*Dxz4* double deletion (deletion on Xa) lack the *Firre* lncRNA, (2) females carrying a paternal *Firre* deletion (deletion on Xi) lack the superloop and (3) females carrying a paternal *Dxz4* single or *Firre*-*Dxz4* double deletion (deletion on Xi) lack both the superloop and megadomains (Fig. 1f left). To identify dysregulated X-linked genes for each possible combination of the deletion, we performed differential expression analysis and found that the only dysregulated gene on the X chromosome was the lncRNA *Firre* (FDR ≤ 0.1 & |log2foldchange| ≥ 1), suggesting that the mega-structures and the *Firre* lncRNA have no impact on imprinted XCI (Fig. 1f right).

To further support this result, we performed an allele-specific expression analysis to test if we detect deviations of the expected maternal ratios or a gain of gene escape in the presence of the deletions. We found that the median allelic ratio of all the X-linked genes was unchanged regardless if the deletions were on Xa or Xi (Fig. 1g). Moreover, deletion of these loci on the Xi did not result in increased gene escape in the placenta (Supplementary Fig. 3a). Random XCI in the brain might also not be affected, since we detect the expected XCI skewing ratios in the presence of the deletions, a well-documented effect in female cells from crosses between Bl6 and CAST that results in the predominant inactivation of the Bl6 X chromosome ^19^(Supplementary Fig. 3b). In contrast, for autosomal genes we detect the expected biallelic ratios across all samples (Supplementary Fig. 3c).

To address whether the absence of both the mega-structures and *Firre* RNA has an impact on gene expression in an organ-specific manner, we generated a transcriptomic bodymap from adult females carrying a homozygous double *Firre*-*Dxz4* deletion (Fig. 2a, Supplementary Fig. 4 a-b). We then performed differential expression analysis of spleen, brain, kidney, heart, lung and liver and found that most of the dysregulated genes show organ-specific expression changes, primarily on autosomes (autosomes: 98.15% n=372 chrX: 1.85% n=7), with only a few overlapping across organs (Fig. 2b, Supplementary table A). The highest number of differentially expressed genes was detected in the spleen (n=239, compared to the rest of the organs that on average had 30 dysregulated genes). To validate and categorize the dysregulated genes identified in the DKO into superloop, megadomain, and *Firre* locus dependent gene sets, we sequenced the spleen and the liver from independently generated *Firre* and *Dxz4* SKO (Fig. 2c, Supplementary Fig. 4 B). We hypothesized that dysregulated gene sets that are: (1) shared across the three strains are superloop specific (2) shared between *Dxz4* and DKO are megadomain-dependent and (3) shared between *Firre* and the DKO are *Firre* locus specific. To identify gene sets for these categories, we selected the genes dysregulated in at least one of the three KO strains and where the fold change direction in either one of the single KO agrees with the direction in the DKO (|log2foldchange| ≥ 1). We first used this approach for the liver, an organ with a low number of differentially expressed genes in the DKO, and identified 15 megadomain, 4 superloop, and 4 *Firre* locus dependent genes (Fig. 2d left, Supplementary table B). Interestingly, we detected megadomain dependent upregulation of an entire gene cluster on chromosome 4 (Fig. 2d right). Notably, without applying the fold change cutoff, we observe megadomain dependence of *Xist*, which is significantly upregulated in the DKO brain and kidney, as well as in the Dxz4 SKO spleen (mean upregulation 30%, Supplementary Fig. 4 c).

**Figure 2.**
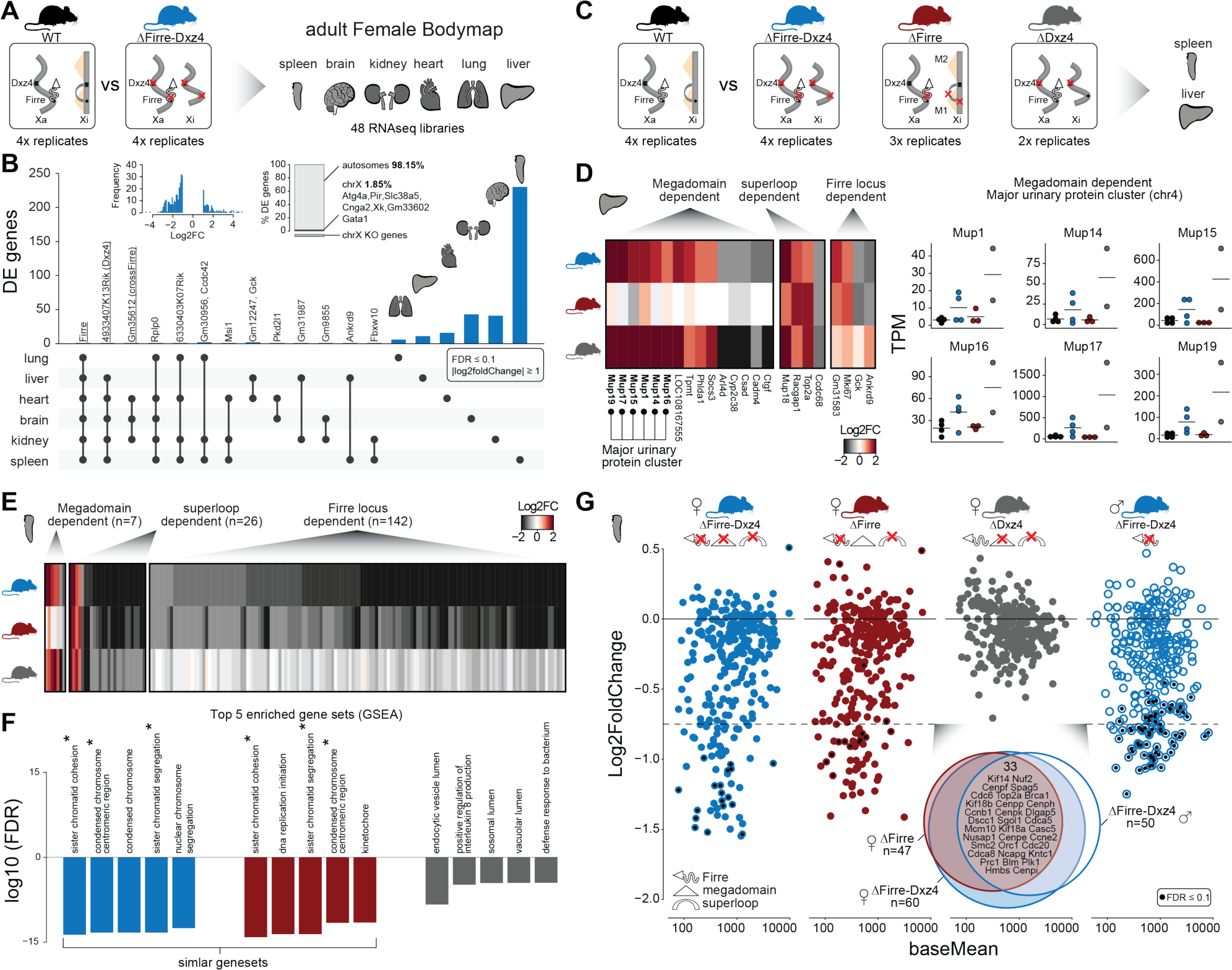
Homozygous deletion of *Firre* and *Dxz4* loci results in organ-specific expression changes on autosomes. (a) Cartoon illustrating the structural differences of the X chromosome between wildtype and double KO female mice (left), and the organs collected from 4 WT and 4 KO six-weeks old adult females to generate the transcriptomic bodymap (right). (b) Overview of the differentially expressed genes in the female bodymap and their overlap across the 6 organs (DEseq2: FDR ≤ 0.1, |log2foldChange| ≥ 1). The histogram displays the log2foldChange distribution of the differentially expressed genes. The bar plot shows the proportion of differentially expressed genes between autosomes and the X chromosome. Genes that are directly affected by the deletion are underlined (*Firre, Dxz4* transcript (*4933407K13Rik*) and *crossFirre* (*Gm35612*, antisense to *Firre*). (c) Cartoon illustrating the structural differences of the X chromosome between WT and each of the KO strains (left), and the SKO organs collected from six-weeks old adult females (Firre=3, Dxz4=2) (right). (d) Heatmap showing mthe fold changes of megadomain, superloop and *Firre* locus dependent gene sets in the liver (left). The “major urinary protein” gene family cluster on chromosome 4 shows megadomain dependent upregulation (right). (e) Heatmap showing the fold changes of megadomain, superloop and *Firre* locus dependent gene sets in the spleen. (f) Top 5 enriched dysregulated gene sets in the spleen of the DKO and of the *Firre* and *Dxz4* SKO (p-value corrected for multiple testing using the p.adjust function in R according to the Benjamini & Hochberg method). The GSEA analysis was performed on DEseq2 test statistics with all GO gene sets (c5.all.v6.2.symbols). Asterix indicates the overlapping gene sets. (g) MA plot for all KO strains of the genes extracted from the top 5 gene sets identified in the DKO and *Firre* SKO. The black dot indicates significant differentially expressed genes. Venn diagram showing the overlap between female and male DKO and *Firre* SKO genes with a log2foldchange ≤ −0.75.

We next applied the same approach to the spleen, the organ with the highest number of differentially expressed genes in the DKO, and identified 7 megadomain (4%), 26 superloop (14.8%) and 142 *Firre* locus (81.1%) dependent genes (Fig. 2e, Supplementary table C). The *Firre* locus dependent gene set, the largest group, contains only downregulated genes. By gene set enrichment analysis (GSEA), we find that the top *Firre* SKO and DKO enriched gene sets are almost identical, sharing downregulated gene sets involved in chromosome structure and segregation (Fig. 2f). Genes extracted from the enriched gene sets identified in the *Firre* SKO and DKO share a similar pattern of downregulation, which was not observed in the *Dxz4* SKO (Fig. 2g). In addition, fold changes in *Firre* SKO and DKO are strongly correlated (spearman rho = 0.86, *P*-value = 2.2*10^-16^), indicating that the *Firre* locus is the main driver for downregulation of these gene sets (Fig. 2g). To test whether this molecular phenotype can be uncoupled form X inactivation, we performed the same analysis in DKO male spleens. We found a similar pattern of downregulation as observed in females (Fig. 2g right) and we also found that across all strains carrying the *Firre* locus deletion, the enriched gene sets share the majority of genes with the greatest degree of dysregulation. Taken together these findings point to the *Firre* locus, independently of X inactivation, as the main driver of these autosomal expression signatures.

## Discussion

The *Firre* and *Dxz4* loci provide the platform for the formation of the X chromosome mega-structures and have been extensively studied in cell lines modeling random XCI ^9–13^. Here we addressed the *in vivo* role of these elements by generating mice carrying a single or double deletion of these loci. In agreement with previous *in vitro* studies, we find that the loss of these loci *in vivo* does not affect random XCI. The lack of dysregulated X-linked genes in adult organs suggests that these mega-structures may also not be important for long-term maintenance of the Xi chromosome. Moreover, by studying the placenta we were able to show for the first time that loss of these loci also does not affect imprinted XCI.

Remarkably, deletion of these loci results in reproducible organ-specific expression changes on autosomes suggesting that structural changes of the Barr body may lead to autosomal gene dysregulation. Indeed, crosstalk between autosomes and the X chromosome has been proposed as a mechanism for X inactivation counting ^20^. Whether these changes are directly regulated by the Barr body structure remains to be investigated. The largest transcriptional effect on autosomes is a *Firre* locus-dependent and X inactivation-independent, suggesting a role for the *Firre* RNA or DNA locus in autosomal gene regulation. However, the finding that the majority of the dysregulated genes are on autosomes and involved in chromosome structure and segregation, which is in line with the known role for *Firre* lncRNA in nuclear organization ^14^, points to an RNA dependent role for autosomal gene regulation. Collectively, our results indicate that the X-linked loci *Firre* and *Dxz4*, are involved in autosomal gene regulation rather than XCI biology *in vivo*.

### Accession codes

Sequence data and alignments have been submitted to the Gene Expression Omnibus (GEO) database under accession code GSE127554.

## ACKNOWLEDGEMENTS

We thank Philipp Maass, Marta Melé, Rasim Bracutu, Kaia Mattioli, Gabrijela Dumbovic and Quanah Hudson for stimulating discussions and critical reading of the manuscript. Sequencing was performed at the Bauer Core Facility at Harvard University.

## AUTHOR CONTRIBUTIONS

D.A., J.L.R., A.M. designed the study. D.A. and C.G. performed the experiments. Z.D.S generated the Dxz4 and Firre-Dxz4 double KO mouse. J.P.L. generated Firre SKO. D.A. performed all the bioinformatic analysis. D.A. and J.L.R. wrote the manuscript with input from the authors. All authors have read and approved the manuscript for publication.

## Materials and methods

### Mouse strains

Mice were housed under controlled pathogen-free conditions (Harvard University’s Biological Research Infrastructure). C57BL/6J (Bl6), B6D2F1/J (F1 Bl6 and DBA) and CAST/Ei (CAST) mice were purchased from the Jackson Laboratory. The *Firre* deletion mouse (deletion mm10: chrX:50,555,286-50,637,116) was generated from genetically modified embryonic stem cells (129xBl6 background) by inserting 2x LoxP sides upstream and downstream of the *Firre* gene body followed by crossing the founder mouse with CMV-CRE (BALB/C background), as described in detail in (Lewandowski et al., unpublished). The *Dxz4* single deletion (chrX: 75,721,164-75,764,733 mm10) and the *Firre*-*Dxz4* double deletion strains (Dxz4 deletion chrX:75,720,836-75,764,839, Firre deletion same as for Firre SKO), were generated by co-injecting Cas9 mRNA together with two guide RNA’s that span the Dxz4 locus into pronuclear stage 3 (PN3) zygotes as previously described ^1^ (supplementary Figure 1a). Zygotes were either isolated after mating superovulated B6D2F1 female mice (Jackson labs) with Bl6 males for the *Dxz4* single KO (SKO) or generated by piezo-assisted Intracytoplasmic sperm injection of *Firre* SKO sperm into B6D2F1 oocytes for the double KO (DKO) (protocol described in ^2^. At PN3, Cas9 mRNA (200 ng/ul) was co-injected with the spanning gRNAs (50 ng/ul each), after which embryos were cultured to the blastocyst stage, transferred into pseudopregnant CD-1 strain females (Charles River), and brought to term. Protospacer sequences (shown in Supplementary Figure 1) were identified using ChopChop ^3^ and single guide RNAs were synthesized from T7 promoter containing oligonucleotides using the MEGAshortscript™ *in vitro* transcription system (Invitrogen). Founder mice from the three strains were backcrossed at least two times with C57BL/6J to remove CRISPR-Cas9 off target effects.

### Tissue isolation and library preparation

To determine whether the deletion of *Firre, Dxz4* or *Firre*-*Dxz4* impact random or imprinted X chromosome inactivation, we collected female and male embryonic day 12.5 brains and placentas from reciprocal F1 crosses between the deletion strains and CAST/EiJ. From every strain and tissue we collected 9 wildtype samples (forward cross: 3 male and 3 female, reverse cross: 3 female) and 9 that carry a deletion on the maternal or paternal allele (forward cross: ΔxCAST 3 male and 3 female, and reverse cross: 3x CASTxΔ, maternal allele always on the left). To reduce the amount of maternal contamination we removed the decidua of the placentas.

For the *Firre*-*Dxz4* adult bodymap, we collected the spleen, brain, kidney, heart, lung and liver from 6 weeks old female mice carrying a homozygous double deletion (4 WT and 4 DKO replicates). To validate and classify the dysregulated genes form the DKO, we collected liver and spleen from independent generated female SKO strains (*Firre* 3 SKO and *Dxz4* 2 SKO replicates). To test whether the molecular phenotype observed in the female spleen can be uncoupled form X inactivation, we performed the same analysis in DKO male spleens (2 WT and 2 DKO replicates).

The collected tissues were snap frozen and stored at −80°C until further processed. RNA was extracted from TRIzol using standard protocols. The Illumina TruSeq kit was used to create polyA^+^ libraries from total RNA. We generated strand-specific libraries for tissues collected form the F1 crosses (TruSeq stranded Illumina) and non-strand-specific TruSeq libraries for the adult organs. All the libraries were run on a Bioanalyzer to assess purity, fragment size, and concentration and sequenced on a HiSeq 2500 at Harvard University’s Bauer Sequencing Core (75 bp paired end).

### Sequencing alignment and analysis

The RNA data was aligned with STAR by using specific parameters to exclude reads mapping to multiple locations (STAR version 2.5.0c: –outFilterMultimapNmax 1) ^4^. The read counts for every isoform within the RefSeq gene annotation (downloaded February 2018) were calculated by using the Python script htseq-count (HTSeq version 0.6.1) ^5^. Differential expression analysis was performed with standard settings of DESeq2 (DESeq2 version 1.22.1, R version 3.5.1).

Allele-specific expression was detected from RNA-seq by using the Allelome.PRO, as described in detail in Andergassen et al. 2015 ^6^. Allelome.PRO uses the information of characterized single-nucleotide polymorphisms (SNPs) to assign sequencing reads to the corresponding strain in F1 crosses. For the SNP annotation we first extracted 20,606,390 high confidence SNPs between the CAST/EiJ (CAST) and C57BL6NJ (Bl6) form the Sanger database as described previously ^6,7^. Although we backcrossed all the founder mice serval times to Bl6, there is a possibility that a few regions in the genome are 129, BALB/C or DBA background (originating from strains that were used to generated the KO founder strains). In order to correct our allele-specific analysis for such regions, we only used Cast/Bl6 SNPs where the Bl6 allele was shared between the strains 129, BALB/C and DBA (Final SNP number: 15,438,314 SNPs). For the Allelome.PRO analysis we only included SNPs that are covered by at least 2 reads in by setting the “minread” parameter to 2.

## Supplementary Figures

**Supplementary Figure 1.**
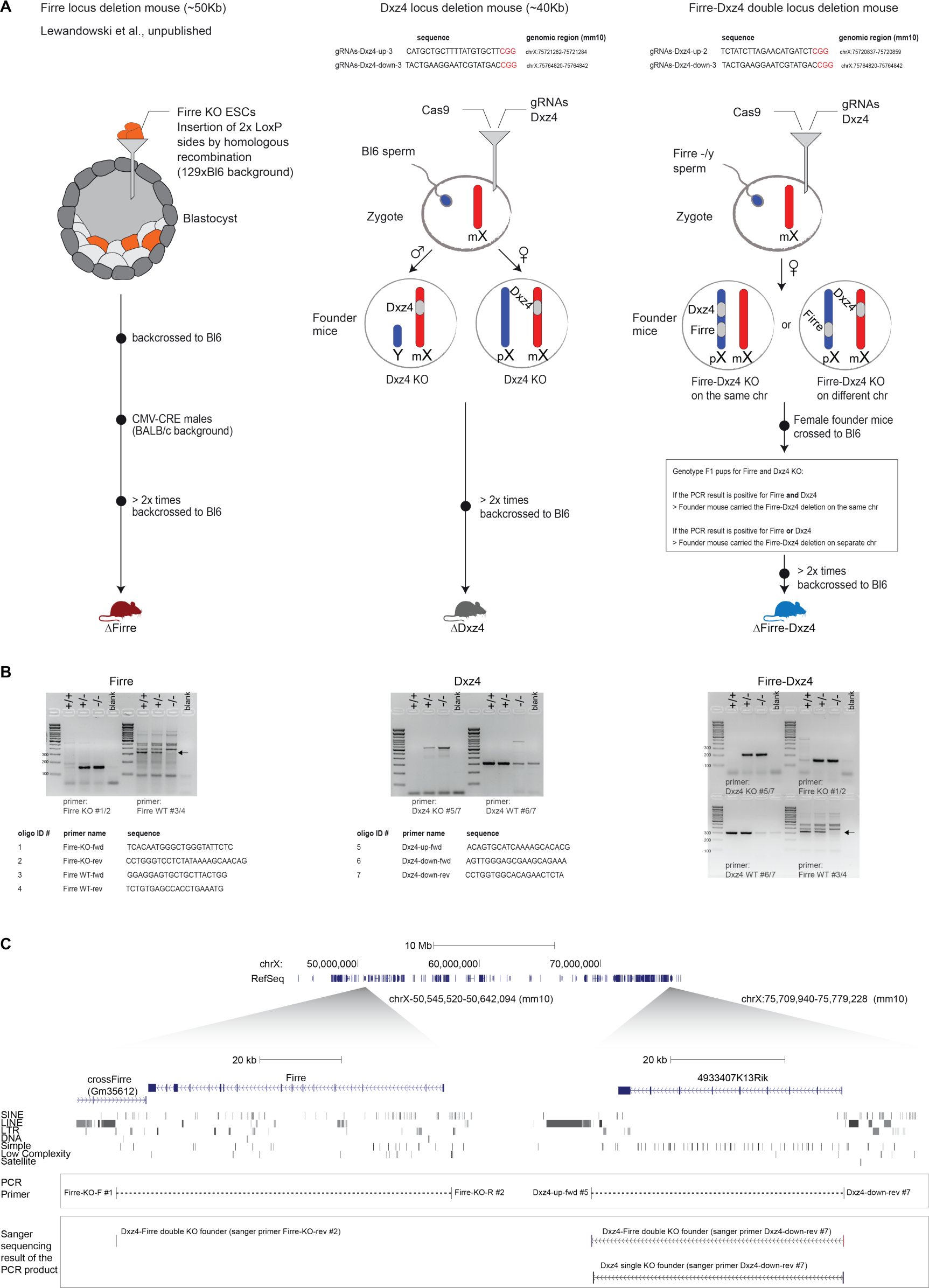
Generation of mice that carry a single or double deletion of *Firre* and *Dxz4*. (a) Schematic overview of the three genome-editing strategies, more details in the material and methods section. (b) Genotype strategy and required primers to identify the KO and WT allele for *Firre* and *Dxz4* (c) UCSC genome browser showing the Firre and Dxz4 region. The PCR product of the KO bands was Sanger sequenced and the resulting sequence was aligned to the UCSC genome browser to confirm the deletion of these loci.

**Supplementary Figure 2.**
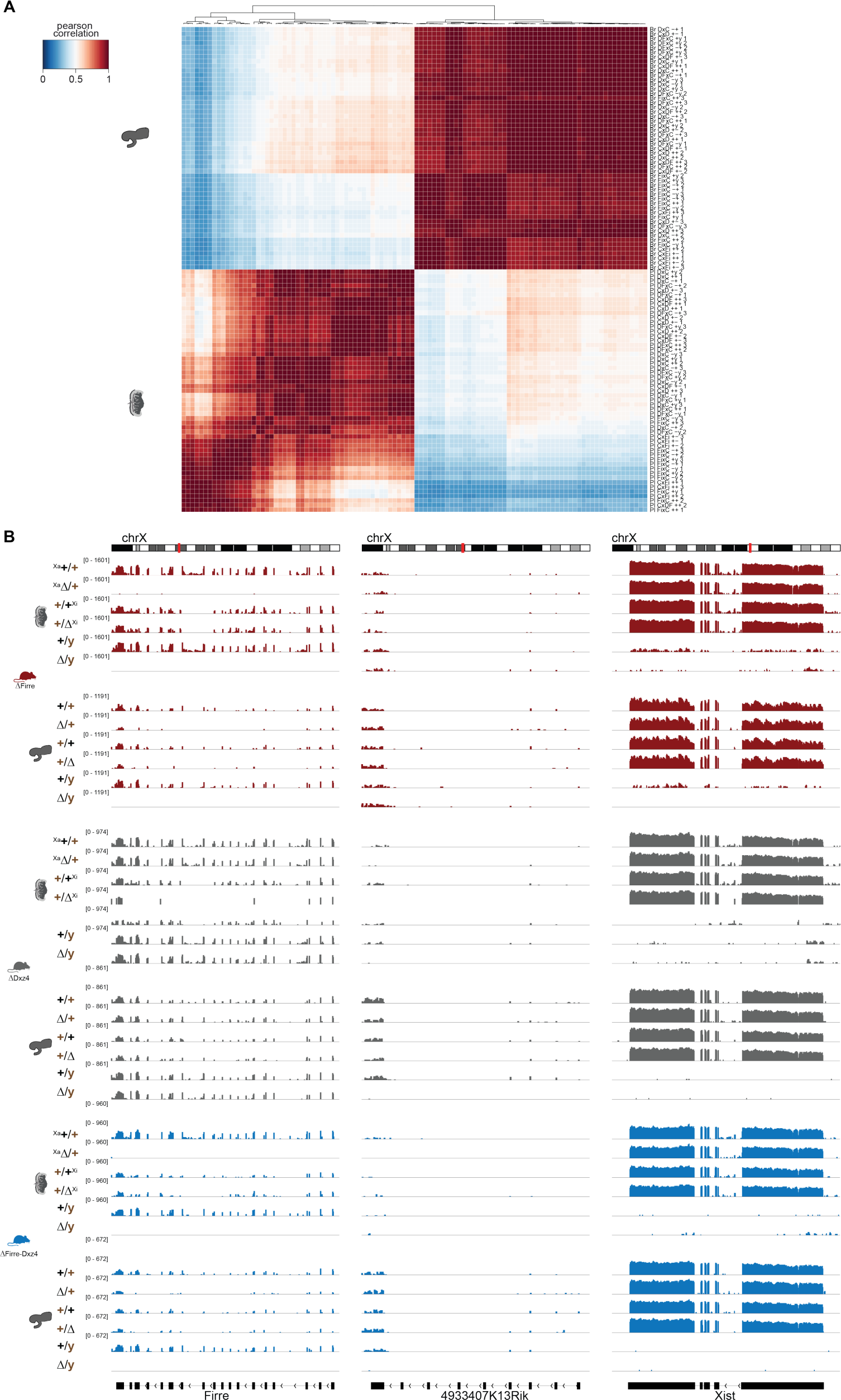
RNA-seq quality control from F1 brains and placentas. (a) Heatmap showing unsupervised clustering of a Pearson correlation matrix (106 placenta and brain samples) from expression data (TPM), confirming the expected developmental relationship. (b) *Firre, Dxz4* (*4933407K13Rik*) and *Xist* expression abundance in placenta and brain collected from the forward (maternal inheritance of the deletion) and reverse cross (paternal inheritance of the deletion) for all three strains. Only one of the replicates is shown.

**Supplementary Figure 3.**
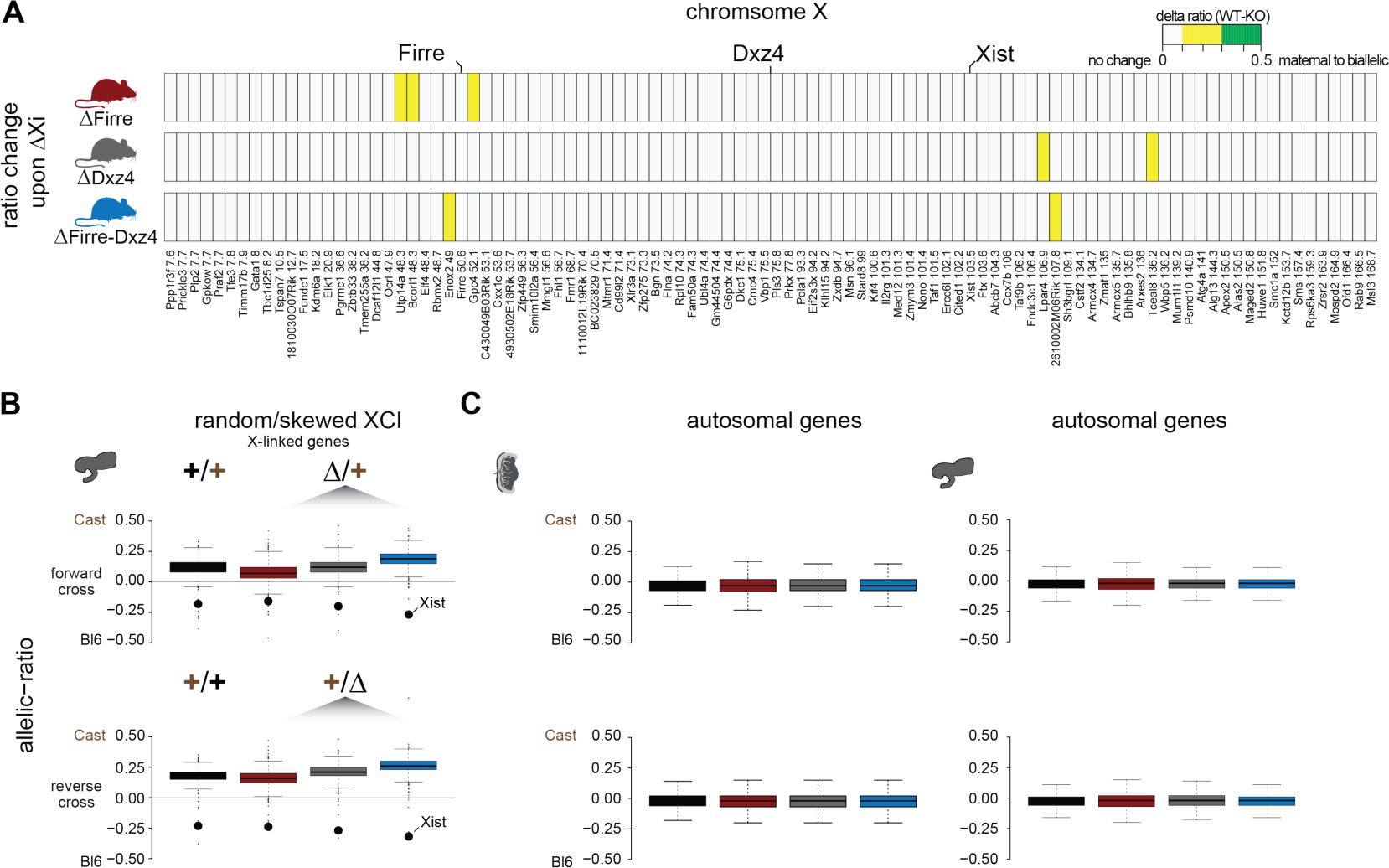
Allelic ratio analysis from F1 brains and placentas. (a) Heatmap showing the ratio change of X-linked genes in the placenta between wildtype and having the deletion on the Xi (paternal X) for the three KO strains. The top 100 genes with the highest deviations are shown. White: ratio change <10%, yellow: 10% ≤ ratio change <30%, green: ratio change ≥ 30%. (b) Boxplot showing the allelic ratio of X-linked genes for the brain, an organ that undergoes random/skewed XCI (Bl6 X chromosome in 70% of cells inactive). Allelic ratios are shown for WT and KO strains in the forward cross (top, deletion on the maternal X) and reverse cross (bottom, deletion on the paternal X). (c) Boxplot showing the allelic ratio on autosomes for WT and the three KO strains in the brain and placenta.

**Supplementary Figure 4.**
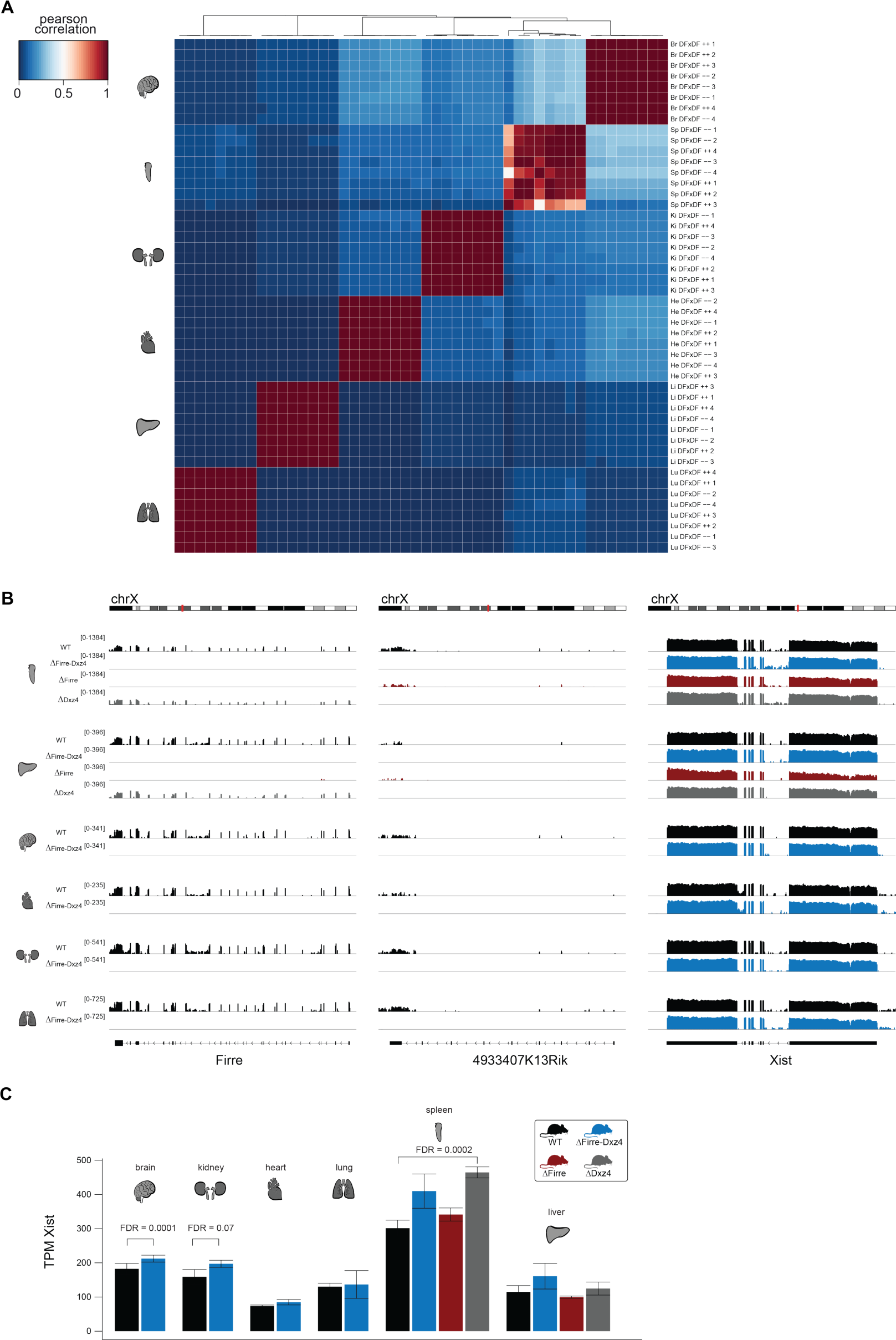
Analyzing adult female organs carrying a homozygous deletion of *Firre* and *Dxz4*. Heatmap showing unsupervised clustering of a Pearson correlation matrix (48 samples, *Firre*-*Dxz4* double deletion bodymap) from expression data (TPM), confirming the expected developmental relationship. The *Firre*-*Dxz4* bodymap includes spleen, brain, kidney, heart, lung and liver from 6 weeks old females (4 WT and 4 KO replicates). *Firre, Dxz4* (4933407K13Rik) and *Xist* expression abundance in the bodymap organs. One of the replicates is shown. (c) Bar plot showing the *Xist* expression abundance in the bodymap organs. FDR values obtained by DEseq2 analysis.

